# Microbial invasion of a toxic medium is facilitated by a resident community but inhibited as the community co-evolves

**DOI:** 10.1101/2022.03.03.482806

**Authors:** Philippe Piccardi, Géraldine Alberti, Jake M. Alexander, Sara Mitri

## Abstract

Predicting whether microbial invaders will colonize an environment is critical for managing natural and engineered ecosystems, and controlling infectious disease. Invaders often face competition by resident microbes. But how invasions play out in communities dominated by facilitative interactions is less clear. We previously showed that growth medium toxicity can promote facilitation between four bacterial species, as species that cannot grow alone rely on others to survive. Following the same logic, here we allowed other bacterial species to invade the four-species community, and found that invaders could more easily colonize a toxic medium when the community was present. In a more benign environment instead, invasive species that could survive alone colonized more successfully when the residents were absent. Next, we asked whether early colonists could exclude future ones through a priority effect, by inoculating the invaders into the resident community only after its members had co-evolved for 44 weeks. Compared to the ancestral community, the co-evolved resident community was more competitive toward invaders, and less affected by them. Our experiments show how communities may assemble by facilitating one another in harsh, sterile environments, but that arriving after community members have co-evolved can limit invasion success.

## Introduction

Successful colonization of invader microorganisms into sterile environments or existing microbial communities are common, and can impact ecosystem diversity and function, potentially with dramatic consequences (1–3). A better understanding of the factors driving microbial invasions may help to prevent the spread and establishment of invasive species, or to aid the intentional introduction of a new species for a desired purpose. For example, it might be desirable to prevent the invasion of a species that reduces the efficiency of a bioremediation system (4), or to promote the colonization of probiotic species in the intestinal microbiome of a patient (5, 6).

What determines the ability of an invasive species to colonize an existing ecosystem depends on the characteristics of both the invading species and the resident community (7, 8). Many theoretical and empirical studies have established factors that influence invasion outcome, such as propagule pressure (9–14), resident community productivity (15), genotypic richness of invaders (13, 16) or the resident community (3, 13, 17–20), community niche coverage (3, 21, 22) and abiotic conditions (e.g. the presence of antibiotics (23)).

Invasion success may also depend on the sign and strength of interactions between resident community members, and between residents and invaders (24). Previous studies tend to find that invaders compete with resident species (3, 13, 17, 20, 23, 25, 26), which is consistent with competition typically dominating microbial communities (27–29). However, in sterile environments, early colonizers can facilitate the arrival of other species (30–35). This occurs when new communities assemble and groups of species follow one another in so-called “successions”, for example in the formation of dental plaque (36, 37) or marine particle communities (32, 33). Facilitation likely occurs in newly assembling communities, as sterile environments are typically difficult to colonize, for example if they have an extreme pH, contain toxic compounds, or are lacking in easily accessible nutrients or water. Pioneer species may alter the environment in ways that facilitate invasion by new species that would otherwise not survive (24, 38–41). This is in line with the Stress Gradient Hypothesis (SGH), which predicts that species are more likely to interact positively in stressful environments (42–48). The link between the SGH and microbial invasion has, however, not yet been experimentally tested.

As more species colonize the environment and species diversity increases, previously available niches begin to fill up, such that competition is expected to increase and invasion success to drop. The negative relationship between invasion success and species richness and diversity have been well-established (3, 13, 17–20). As time passes, resident species may co-evolve to reduce niche overlap and availability in a way that would prevent further invasion. The Community Monopolization Hypothesis predicts that early colonisers adapt to use available resources efficiently, yielding a competitive advantage against later-arriving species (49–52), also known as a “priority effect” (31, 53). One may also expect such co-evolved resident species to be less perturbed by species invasion (49). Experimentally disentangling the role of the different factors discussed above on invasion success and resistance can be challenging.

Here we aim to test the effect of the two less-well understood factors (the SGH and priority effects) on bacterial invasion success by studying invasion into a synthetic bacterial community whose composition is fixed at four species: *Agrobacterium tumefaciens, Comamonas testosteroni, Microbacterium saperdae* and *Ochrobactrum anthropi*. These four species can grow and bioremediate Metal Working Fluids (MWF) (47, 54), an industrial fluid used in metal manufacturing. MWFs contain mineral oils, emulsifiers and biocides, some of which are toxic to bacteria. In previous work (47), we showed that when the four species were grown together in this toxic environment, they facilitated each other’s survival compared to when they were alone. Instead, when we added amino acids to make the environment more permissive, competition between species increased. This system allows us to study biological invasion while experimentally manipulating environmental conditions to control interactions between community members and holding all other factors constant. Another advantage of this system is that it stabilizes over evolutionary time-scales, allowing us to explore the effect of community co-evolution on microbial invasion.

Using four invader species, *Aeromonas caviae, Klebsiella pneumoniae, Providencia rettgeri*, and *Pseudomonas fulva* that were isolated from waste MWF (chosen from a set of 20 based on our ability to distinguish them from the resident species), we first show that the resident community facilitates invasion of species that cannot grow alone, but inhibits those that can. Whether or not species could grow alone was modulated by changes in the growth medium, in particular the addition of amino acids as nutrient supply. Second, after co-evolving the four resident species for 44 weeks, we found that invasions were still possible in MWF, but the growth of the invaders was inhibited relative to the ancestral community and the co-evolved resident species were less affected by invasions. Together, our results show that facilitative communities are easier to invade than competitive ones, but that a co-evolved community is more robust to invasion compared to an ancestral one.

## Results

### The resident community facilitates the invasion of species that cannot grow alone

We first ask whether the four invader species (*A. caviae, K. pneumoniae, P. rettgeri*, and *P. fulva*), could individually colonize MWF and to what extent the resident community promotes or inhibits invasion. The resident community was cultured in MWF for 1 week, at the end of which 1% of the population was transferred into fresh media, and this was repeated for a total of 4 weeks. Each invader species was inoculated into three replicate microcosms of the resident community 48 hours after the first transfer, presumably during the community’s exponential growth phase (Fig. S1A). The resident community was always invaded by a single invader species at a time. As a control treatment, we inoculated each of the invader species into sterile MWF and performed transfers in parallel (Fig. S1B). The abundance of the invading and resident species was quantified at inoculation and before each transfer (invader species in Fig. 1A; resident species in Fig. S2A, left).

**Fig. 1.**
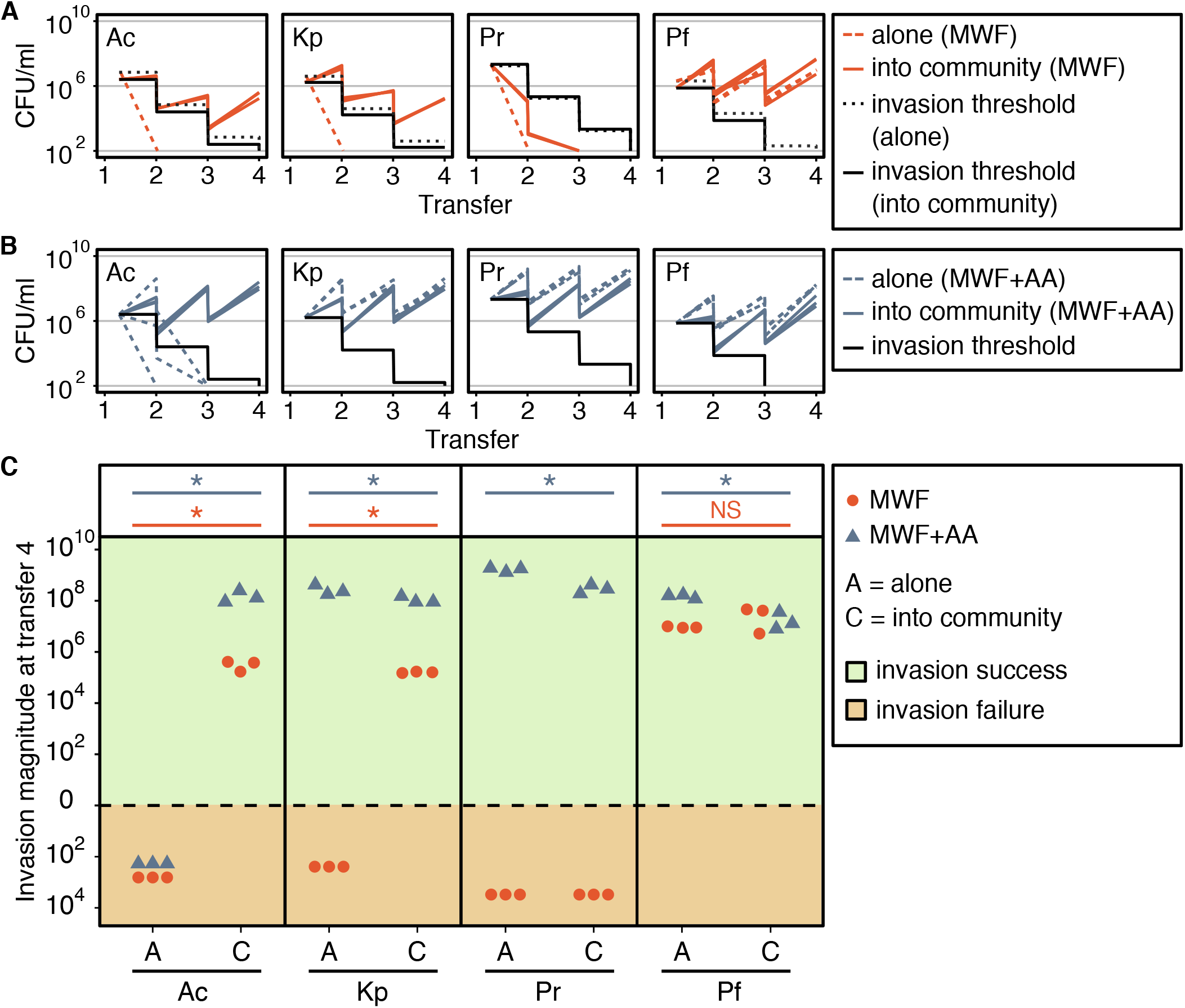
Outcome of invasion in MWF or MWF+AA. (A) Abundances of the four invader species in MWF at different transfers, quantified at first inoculation and before each transfer. The four invader species were grown alone or inoculated into the growing resident community after it had been transferred once and until transfer 4 (1 transfer every 7 days). At each transfer, the culture is diluted 100-fold. The experiments were done in parallel except for the invader alone, which was done separately. The black dotted line and the black solid line represent the invasion thresholds of the four invader species alone and into the community, respectively, which represents the abundance of a non-growing invader over time. We count an invasion as successful if the abundance of the invader at the end of the transfers is higher than the invasion threshold. (B) Abundances of the four invader species in MWF+AA at different transfers. The black solid line represents the invasion threshold when grown alone “A” or invaded into the community “C”. The experiments were all conducted in parallel. (C) To exclude the effect of the choice of dilution rate on invasion success, we calculate the “invasion magnitude”: the abundance of each invader at transfer 4 minus its corresponding invasion threshold. If this number is higher or lower than zero, then the invasion is successful (+) or failed (-), respectively. From left to right: *A. caviae* (Ac), *K. pneumoniae* (Kp), *P. rettgeri* (Pr), *P. fulva* (Pf). Statistical significance is marked above the data points (P-values: * <0.05, NS = not significant). For abundance of resident species see Fig. S2.

*P. fulva* was the only one of the four invader species that could colonize the MWF when alone (Fig. 1A (red dotted line), C). Instead, when the community was present, the number of successful invasions increased: *A. caviae*, *K. pneumoniae* and *P. fulva* colonized the MWF containing the resident community, while *P. rettgeri* still did not (Fig. 1A (red solid line), C). These results are in line with our previous findings (47) that species that cannot grow alone in MWF are likely to be facilitated by other species, explaining why in some cases invasion is only successful when the community is present.

### Residents inhibit invaders that can grow alone in a more permissive medium

MWFs are designed to prevent bacterial contamination and are therefore quite toxic. This explains why only one of the invader species was able to grow alone in the MWF medium. To explore invasion in a less harsh environment, we enriched the medium by adding 1% casamino acids (MWF+AA). Casamino acids are a nutrient source for 3 out of the 4 resident community members and, according to previous work (47, 55), we expect more negative interactions in a more permissive medium. We found that *K. pneumoniae*, *P. fulva* and *P. rettgeri* could colonize MWF+AA alone, while *A. caviae* still suffered from the environmental toxicity (invader species Fig. 1B, C; resident species Fig. S2A, right). The three species that were able to colonize alone were still able to invade the community, but significantly less well compared to when the community was absent (Kruskal-Wallis, all p-values < 0.05, Fig. 1C). Consistent with previous work (47), our results suggest that in this more permissive environment, the community competes with the invaders.

### A resident community co-evolved in MWF is more competitive toward invaders

The capacity to colonize a resident community might depend on community history: resident species that have adapted to one another in a given environment may be more likely to exclude future colonists through a priority effect (49–51). To test this hypothesis, we extend the pre-invasion phase to 44 weeks, allowing the four resident species to adapt to MWF and to each other (see Material and Methods). Next, we mixed one co-evolved isolate of each species and call this the “co-evolved community” (Fig. S1C).

We now ask to what extent the invader species can colonize the co-evolved resident community compared to the ancestral one. We found that while *P. rettgeri* could colonize neither, *A. caviae*, *K. pneumoniae* and *P. fulva* colonized both the ancestral and co-evolved communities (invader species Fig. 2A, B; resident species Fig. S2B). However, all three invader species had a smaller invasion magnitude in the co-evolved compared to the ancestral community (Kurskal-Wallis, *A. caviae* p-value < 0.0005, *K. pneumoniae* p-value < 0.0005, *P. fulva* p-value < 0.05, Fig. 2B). The invasion outcome for *A. caviae* was initially inconclusive, where in one out of two biological replicates the invader went extinct when inoculated into the ancestral community (Fig. S3A, B). We tested whether this was due to variability in propagule pressure (13), but found no evidence for this, as different invasion population sizes of *A. caviae* all converged to a similar population size at transfer 4 (invader species Fig. S4; resident species Fig. S5). We therefore concluded that the death of *A. caviae* in one biological replicate might have been due to a technical error (Fig. S3).

**Fig. 2.**
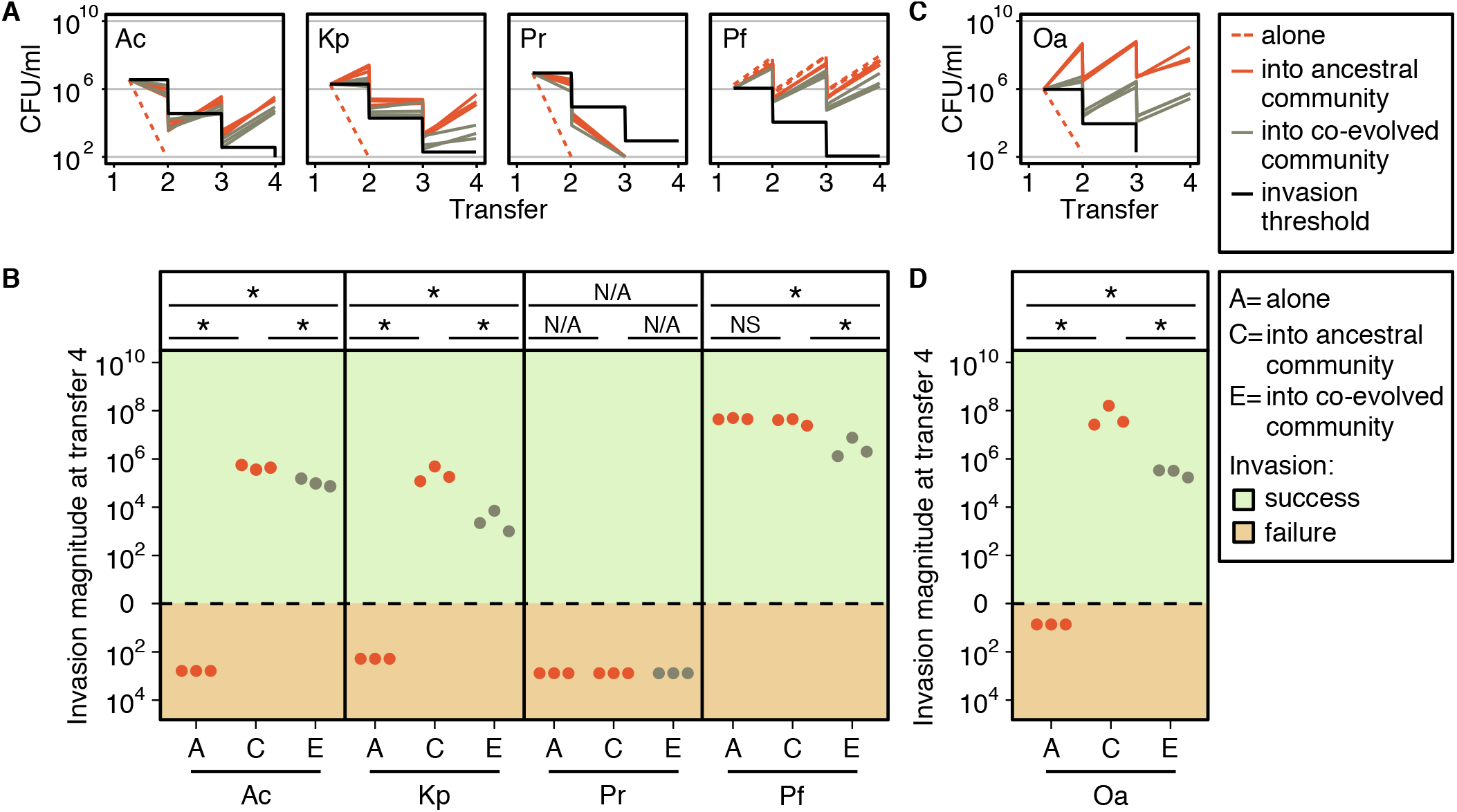
Invasion into ancestral or co-evolved communities. (A, C) Invader species were grown alone, or inoculated into the ancestral or the co-evolved community in MWF. Cultures were diluted 100-fold in fresh MWF every 7 days for a total of 4 transfers. The experiments were all conducted in parallel. The black line represents the invasion threshold. (B, D) Invasion magnitude (abundance at transfer 4 minus the invasion threshold) alone “A”, into the ancestral community “C” or into the evolved community “E”. Positive or negative invasion magnitudes indicate successful (+) or failed (-) invasions, respectively. From left to right: *A. caviae* (Ac), *K. pneumoniae* (Kp), *P. rettgeri* (Pr), *P fulva* (Pf), *O. anthropi* (Oa). Statistical significances are marked above the data points (P-values: * <0.05, NS = not significant, N/A = not applicable statistic). See Fig. 1 caption for more details. In panels A and B, the resident community consists of a co-culture of four ancestral or evolved clonal populations, while in panels C and D, the community contained only three clonal populations (four residents, without *O. anthropi*), either ancestral or evolved. For abundance of the four-species or three-species resident communities see Fig. S2B or Fig. S6, respectively.

We next wondered whether the pattern observed for *A. caviae*, *K. pneumoniae* and *P. fulva* was specific to these invader species colonizing our community of four resident species. One way to explore this is to exclude one species from the co-evolutionary process and allow it to invade at a later stage. We did this by co-evolving three of the resident species, *A. tumefaciens*, *C. testosteroni* and *M. saperdae* together in MWF for 44 weeks, excluding *O. anthropi*. Next, we combined single isolates of the three co-evolved species and invaded the wild-type *O. anthropi* into this co-evolved 3-species community (Fig. S1D). As before, *O. anthropi* could not colonize the MWF when alone (as in (47), Fig. 2C), but invaded successfully when inoculated into the ancestral or the co-evolved community of three. Consistent with the previous invasion assays (Fig. 2A, B), and our hypothesis that a coevolved community is more difficult to invade, *O. anthropi* grew significantly worse when it was inoculated into the co-evolved 3-species community compared to the corresponding ancestral one (invader *O. anthropi* Fig. 2C, D; resident species Fig. S6).

In sum, while invasions into a community co-evolved in MWF are still possible, co-evolved community members inhibit invading species more than their ancestors.

### Co-evolved communities are less affected by invasion compared to their ancestors

So far, we have focused on the effect of the resident community on the invading species. Next, we consider how robust the resident community is to these invasion events.

The abundance of each community member was quantified at the beginning of the experiment and before each transfer. Here, we focus on their abundance at transfer 4, representing cumulative effects (but see e.g. Fig. S2 for remaining data). In all our treatments, the four resident species were maintained over the four transfers. The abundance of two of the ancestral resident species, *A. tumefaciens* and O. *anthropi,* was significantly lower when invaded by *P. fulva* (t-test, both p-values < 0.005, Fig. 3A, Fig. S7A). Otherwise, we detected no significant changes in their abundance following invasion by other species. This lack of perturbation was also observed for *C. testosteroni. M. saperdae’s* abundance was instead greater in the presence of most invaders (t-test, *A. caviae* and *P. rettgeri* both p-values < 0.005, *P fulva* p-value < 0.005, Fig. 3A, Fig. S7A). This is not surprising, as we know that *M. saperdae* strongly depends on other species to grow in MWF (47).

**Fig. 3.**
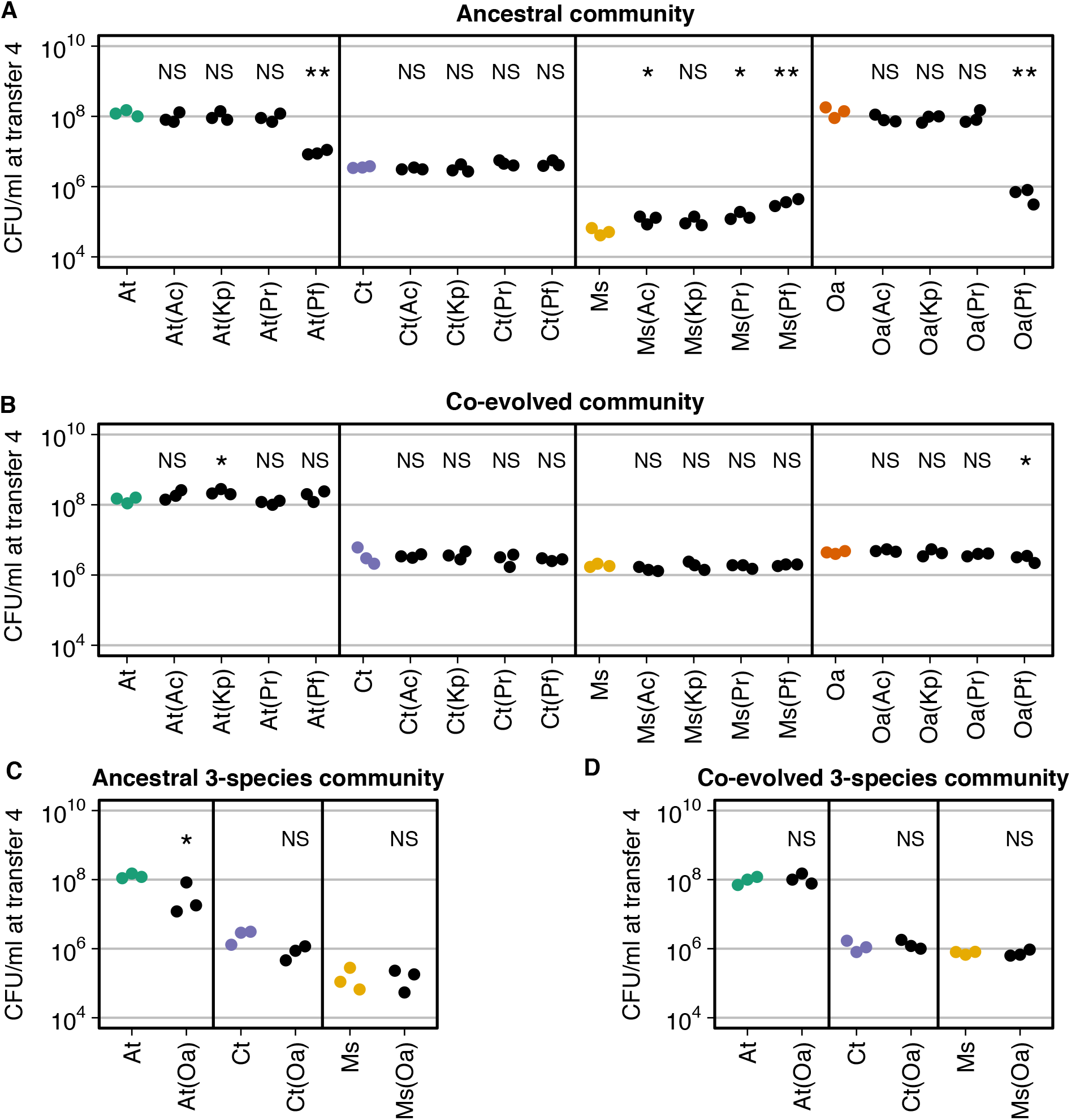
Bacterial abundance of ancestral or co-evolved community members with or without invader. Each panel represents the total population size (CFU/mL) at transfer 4 of a resident member in the ancestral community (A), the co-evolved community (B), the ancestral 3-species community (C) and the co-evolved 3-species community (D). The full datasets are in Fig. S2 and S6. All experiments were performed in MWF. The bacterial abundance of community members without any invader species is represented by colored dots, and once invaded by black dots (invader species indicated in brackets). From left to right: *A. tumefaciens* (At), *C. testosteroni* (Ct), *M. saperdae* (Ms), *O. anthropi* (Oa), *A. caviae* (Ac), *K. pneumoniae* (Kp), *P. rettgeri* (Pr), *P. fulva* (Pf). We compared the data points of each species when invaded to the corresponding data points when co-cultured with other community members but without invasion. Statistical significance is marked above the data points (P-values: * <0.05, **<0.01, NS = not significant).

Once the community had co-evolved, the abundances of *A. tumefaciens* and *M. saperdae* were no longer significantly affected by the invasion of *P. fulva* (Fig. 3B). The abundance of *O. anthropi* was still lower following invasion by *P. fulva*, but significantly less compared to the ancestor (ancestor versus co-evolved, t-test, p-value = 0.0167, Fig. 3A, B, last column). In addition, the abundance of *M. saperdae* was no longer significantly positively affected by any of the invaders. This may be because the co-evolved *M. saperdae* grows significantly better within the resident community (Fig. 3A, B). The co-evolved 3-species community behaved similarly: While the abundance of ancestral *A. tumefaciens* was significantly lower following the invasion of *O. anthropi* (t-test, p-value < 0.05), its co-evolved counterpart was not (Fig. 3C, D). Altogether, co-evolved resident communities were more robust to invasion compared to ancestral ones.

## Discussion

Studies on microbial invasion often focus on how resident community composition and species richness affect invasion outcomes. Here, we chose instead to work with a resident community whose composition was fixed at the same four species, and ask how their environment - specifically environmental harshness - and their common evolutionary history affect invasion success.

By increasing the permissiveness of a harsh medium (MWF) through the addition of amino acids (MWF+AA), the number of invader species able to grow alone increased from a single one to 3 out of 4 species. In almost all cases where invaders died alone, the resident community facilitated their survival and growth. Instead, if invaders could survive alone in the more permissive environment, they experienced a net negative effect if the community was present. This is consistent with our previous research (47), and more generally with the Stress Gradient Hypothesis (SGH), which is rarely linked to invasion ecology. Our results highlight this intuitive link: in a harsh environment colonized by few species, invasion success may be high, as niches are still available and invaders can rely on the presence of the residents to survive. This is expected if early-arriving species improve the environment, facilitating the growth of others that are less well adapted to it (38–41). Alternative dynamics are possible if the first colonizers alter the environment in a way that strongly inhibits future invaders (31), or if late colonizers out-compete earlier ones and replace them (56), but this is not what we observe in our system.

If this intuition is correct, it would explain why the assembly of many natural microbial communities is often highly predictable (32, 33, 36, 37, 57). For example, microbial colonization of the healthy mammalian gut displays specific patterns of species arrival (57, 58). But is a sterile gut a harsh environment? Microbes colonizing a newborn gut must survive the acidic conditions of the stomach, the host’s immune system and bile acids and cholesterol produced by the host that are toxic for most microbial species (59). A few specialized *Lactobacillus* and *Bifidobacterium* species produce bile resistance proteins (60), which allow them to colonize the gut and may facilitate the arrival of other species (61). Similar dynamics may occur in other systems where strong ecosystem perturbations clear the ground for new communities to assemble, such as following antibiotic treatments, or the heavy pollution of soils. However, it remains to be seen whether first colonizers facilitate future arrivals as would be predicted by the SGH (42, 47, 62), or whether it is more of a race to fill available niches.

Once a community has assembled despite the challenging environment, we next asked whether the timing of invasion matters. In our experiments, early invaders fared better than those colonizing a community whose species had co-evolved. But what makes the co-evolved MWF community more resistant and robust to invasion? We noticed that during the last of the 44 transfers, there were relatively small population size fluctuations compared to the first transfers (Fig. S8A, B), suggesting that ecological dynamics had stabilized, which is expected based on similar studies (63). Community stability is often defined as the capacity of a community to return to its initial state after perturbation, such as an invasion (64–67). It is then perhaps not surprising that ecological stability correlated with invasion resistance and robustness (Fig. 3A, B or C, D).

The higher stability - or the higher invasion resistance - of co-evolved communities can be due to several reasons. First, co-evolved *A. tumefaciens* and *M. saperdae* (but not *C. testosteroni* and *O. anthropi*) grew to larger population sizes during the first days (Fig. S9), suggesting that they may take up nutrients faster, making them competitive against invaders. Second, communities with high strain-level diversity tend to be more productive and more robust against invasions (15, 68). However, strain-level diversity and community productivity cannot explain invasion outcomes in our system: by using single isolates from the co-evolved communities, our experiments had the same strain-level diversity in all treatments (Fig. S1A, C); we also observed no significant differences in productivity between ancestral and co-evolved communities (Fig. S10). Third, it could be that the residents co-evolved to actively inhibit other species. If this were true, we might expect the resident species to interact negatively with one another, but on measuring pair-wise interactions between ancestral (Fig. S11G) and co-evolved species (Fig. S12G, Fig. S13F) we only observed that positive interactions weakened between *A. tumefaciens* and *C. testosteroni*, and increased in the other two species (Fig. S12G, Fig. S13G), making this scenario less plausible. Finally, the co-evolving residents may have partitioned the available niches among themselves, leaving little “space” for new arrivals (31). Further investigation, possibly using metabolomics analyses, would be needed to clarify whether this is the case and to more mechanistically understand microbial resistance against invasion.

Our fixed-species experimental design revealed some interesting patterns of invasion success, but also has its limitations. One confounding factor is that adding amino acids to the growth medium allowed more species to grow alone, but also provided new niches for invader species to occupy. This is reflected in the higher overall invasion magnitude of species in MWF+AA compared to MWF (Fig. 1C). But despite these additional niches, invasion magnitude was still lower when the community was present compared to its absence (Fig. 1C). This made it difficult to interpret how invaders affected the resident community grown in MWF+AA: the effects varied depending on the invader species and the resident species with no clear pattern (Fig. S7). Digging deeper into the mechanisms behind the interactions in our system, and developing a theoretical basis for what to expect may help to understand these effects.

One could also question whether a small synthetic community is representative of natural communities and their diversity. A mathematical model in our previous study indicated that competition would increase with a higher number of species in MWF (47), and perhaps we would expect invasions to be less successful in this context. This would also align with experiments involving larger communities that presumably occupy more niches and leave fewer resources for the invader (13, 17–20, 25). Nevertheless, our community could help to understand the first phases of community assembly, when only few species have colonized.

Another weakness of our study is the arbitrary choice to perform four transfers at a 1% dilution rate. To compensate, we were careful to define our measures independently of these choices, such that we could compare between treatments rather than considering absolute measures of invasion. We quantified “invasion success”, representing absolute population increase or decrease, and “invasion magnitude”, which compares population sizes between treatments at the end of the experiment. Another possibility would have been to extend the length of the experiment to observe whether invaders eventually went extinct or established themselves. However, as we were interested in the ecological dynamics of invasion separately from the evolutionary dynamics of the resident community, we decided to keep the invasion time-scale short and assume that species’ genetic adaptation to the environment and each other was negligible. In reality, of course, invaders might acquire mutations that increase invasion success.

In conclusion, we used a model system to disentangle interactions between species and measure their effect on microbial invasion. This revealed that a small, facilitative resident community can improve the environment for species that would otherwise be unable to colonise. However, a community whose residents have adapted to the environment and each other is more difficult to invade. Our work links invasion ecology with the SGH, priority effects and historical contingency (42, 49, 69). We provide a fresh perspective on community assembly as a sequence of invasion events into a harsh environment, where facilitation may be dominant at first as species complement each other, but decreases as niches are occupied through co-evolution.

## Materials and Methods

### Study system

The 4 bacterial species used to assemble the resident community were isolated from MWF (54), and are referred to as: *Agrobacterium tumefaciens* str. MWF001, *Comamonas testosteroni* str. MWF001, *Microbacterium saperdae* str. MWF001 and *Ochrobactrum anthropi* str. MWF001 (as in (47)). The additional four bacterial species, used to invade the resident community, were kindly donated by Peter Küenzi from Blaser Swisslube AG, Hasle-Rüegsau, and we refer to them as: *Aeromonas caviae* str. Blaser001, *Klebsiella pneumoniae* str. Blaser001, *Providencia rettgeri* str. Blaser001, and *Pseudomonas fulva* str. Blaser001. The Metal-Working Fluid (MWF, Castrol Hysol XF, acquired in 2016) was prepared at a concentration of 0.5% (v/v), diluted in water with the addition of selected salts and metal traces to support bacterial growth. We also used MWF medium supplemented with 1% Casamino Acids (Difco, UK) (MWF+AA). All medium compositions are listed in Dataset 1 and are identical to those used in (47).

### Experimental setup

To assemble the resident community, a single isolated colony of each species was selected and inoculated in 10mL of Tryptic Soy Broth (TSB) in Erlenmeyer flasks (50mL), then incubated overnight at 28°C (200 rpm). To achieve exponentially growing bacteria, with a final concentration of ~10^6^-10^7^ CFU/mL, each bacterial species was inoculated at an OD_600_ of 0.05 measured by spectrophotometry (Ultrospec 10, Amersham Biosciences), in 20mL of TSB in Erlenmeyer flasks (100mL) and cultivated at 28°C, shaken at 200 rpm. After 3h, 200*μ*L of each of the four resident species were combined and centrifuged (5 minutes, 10’000 rcf). The bacterial pellet was resuspended in 30mL of MWF or MWF+AA into borosilicate glass tubes (16×125mm, 30mL).

### Transfers

All communities (the 4-species or the 3-species resident communities) were incubated at 28°C and shaken at 200 rpm for 7 days in either MWF or MWF+AA medium. Every week, 300*μ*L (1%) of the week-old culture was transferred into fresh medium and the growth cycle repeated. Each week, we also harvested 1ml of each culture, spun it down at 10,000 rcf for 5 minutes, re-suspended it in glycerol 25% (diluted in PBS) and stocked it at −80°C for future analyses. This was repeated for 44 transfers (weeks) to co-evolve the resident communities or for 4 transfers in the invasion assays. The evolutionary experiment was conducted in 5 replicate culture tubes for each condition (3- or 4-species community), of which we shown only one here (Fig. S8). After the 44 weeks, we isolated one colony of each species, which we refer to as *A. tumefaciens* str. MWF431, *C. testosteroni* str. MWF431, *M. saperdae* str. MWF431 and *O. anthropi* str. MWF431 for the four-species co-evolved community; and *A. tumefaciens* str. MWF351, *C. testosteroni* str. MWF351, *M. saperdae* str. MWF351 and *O. anthropi* str. MWF351 for the 3-species co-evolved community.

### Invasion assays

Invasion was performed after 2 days of the first transfer of the resident community. One single colony of each invader species was selected and inoculated in 10mL of Tryptic Soy Broth (TSB) in Erlenmeyer flasks (50mL) and incubated overnight at 28°C, shaken at 200 rpm. To achieve exponentially growing bacteria, with a final concentration of ~10^6^-10^7^ CFU/mL, each invader strain was inoculated at an OD_600_ of 0.05 measured by spectrophotometry (Ultrospec 10, Amersham Biosciences), in 20mL of TSB in Erlenmeyer flasks (100mL) and cultivated at 28°C (200 rpm). After 3h, 200*μ*L of the invader species were centrifuged (5 minutes, 10’000 rcf). The bacterial pellet was resuspended into the same medium of the resident community. 200*μ*L of this suspension were then added to the culture tubes, with or without the resident communities.

### Bacterial abundance quantification

The abundance of each resident or invader species was quantified before the inoculation in the MWF or MWF+AA (before combining resident species), and before each transfer using serial dilution and selective plating (Fig. 4). To define invasion outcomes we used an invasion threshold representing the dynamics of an invader species with a growth rate of 0 (its abundance changes only due to dilution, i.e. 100-fold decrease every transfer from the initial population size). By subtracting this threshold value from the abundance of the invader species at transfer 4, the invasion is defined as successful if ≤0 (the growth rate is positive) or failed if ≤ 0 (the growth rate is 0 or negative). We used a Kruskal-Wallis test to assess whether effects were significant. We did not used corrections. Why? Raw CFU/mL data and the results of all statistical tests are listed in Dataset 1.

**Figure. 4.**
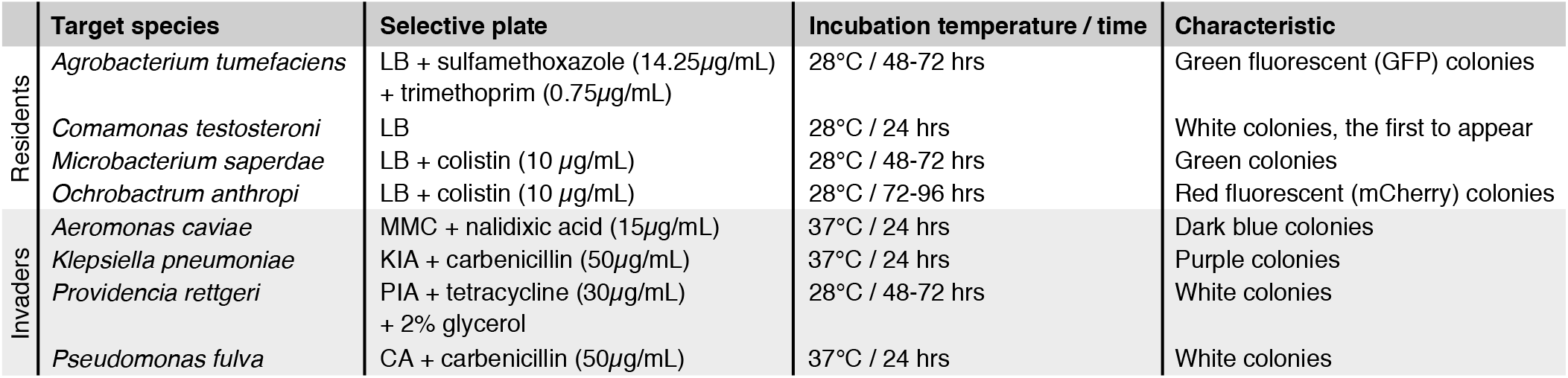
Selective plates list

## ACKNOWLEDGEMENTS

We thank Oliver Meacock and Shota Shibasaki for useful and constructive feedback on the manuscript. We thank Christopher van der Gast and Ian Thompson for providing the 4 species used to assemble the resident community, and Peter Kuenzi for providing the 4 additional species used as invaders. P.P. and G.A. were funded by the University of Lausanne, S.M. by European Research Council Starting Grant 715097. J.M.A. received funding from the European Union’s Horizon 2020 research and innovation programme under grant agreement No. 678841.

## Notes

### Competing Interest Statement

The authors have declared no competing interest.

