## Supplementary material for "Microbial invasion of a toxic medium is facilitated by a resident community but inhibited as the community co-evolves"

### Supplementary figures

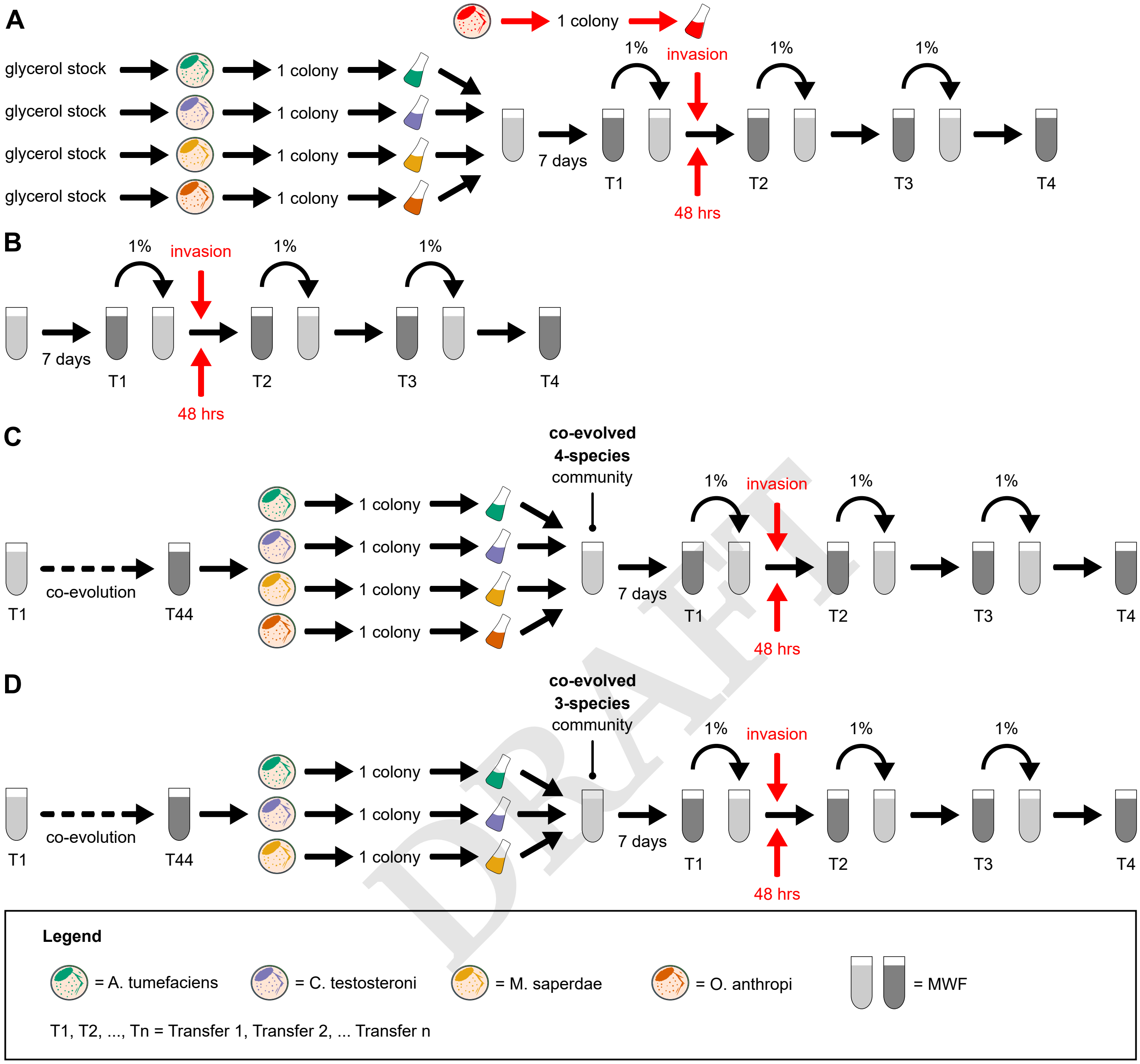


DRAFT

**Fig. S1.** Experimental design. (A) One evolved isolate of each resident species (*A. tumefaciens*, *C. testosteroni*, *M. saperdae* and *O. anthropi*) was randomly picked and grown alone in TSB for 3 hours to exponential phase, 200$\mu$L of each mixed, then washed and resuspended and in 30mL of MWF or MWF+AA. The resident community was cultured in MWF for 1 week, at the end of which 1% of the population was transferred into fresh media, and this was repeated for a total of 4 weeks. In parallel, one colony of the invader species was randomly picked and grown alone in TSB for 3 hours to exponential phase, then washed and 200$\mu$L inoculated into three replicate microcosms of the resident community 48 hours after the first transfer. (B) As a control treatment, the invader species was inoculated into 30mL of sterile MWF or MWF+AA. (C) The four resident species (*A. tumefaciens*, *C. testosteroni*, *M. saperdae* and *O. anthropi*) were co-evolved for 44 transfers. Next, the co-evolved 4-species community was assembled by mixing one co-evolved isolate of each species as in panel (A). Invasion was performed 48 hours after the first transfer. (D) Three resident species (*A. tumefaciens*, *C. testosteroni* and *M. saperdae*) were co-evolved for 44 transfers. Next, the co-evolved 3-species community was assembled by mixing one co-evolved isolate of each species as in panel (A). Invasion was performed by *O. anthropi* 48 hours after the first transfer.

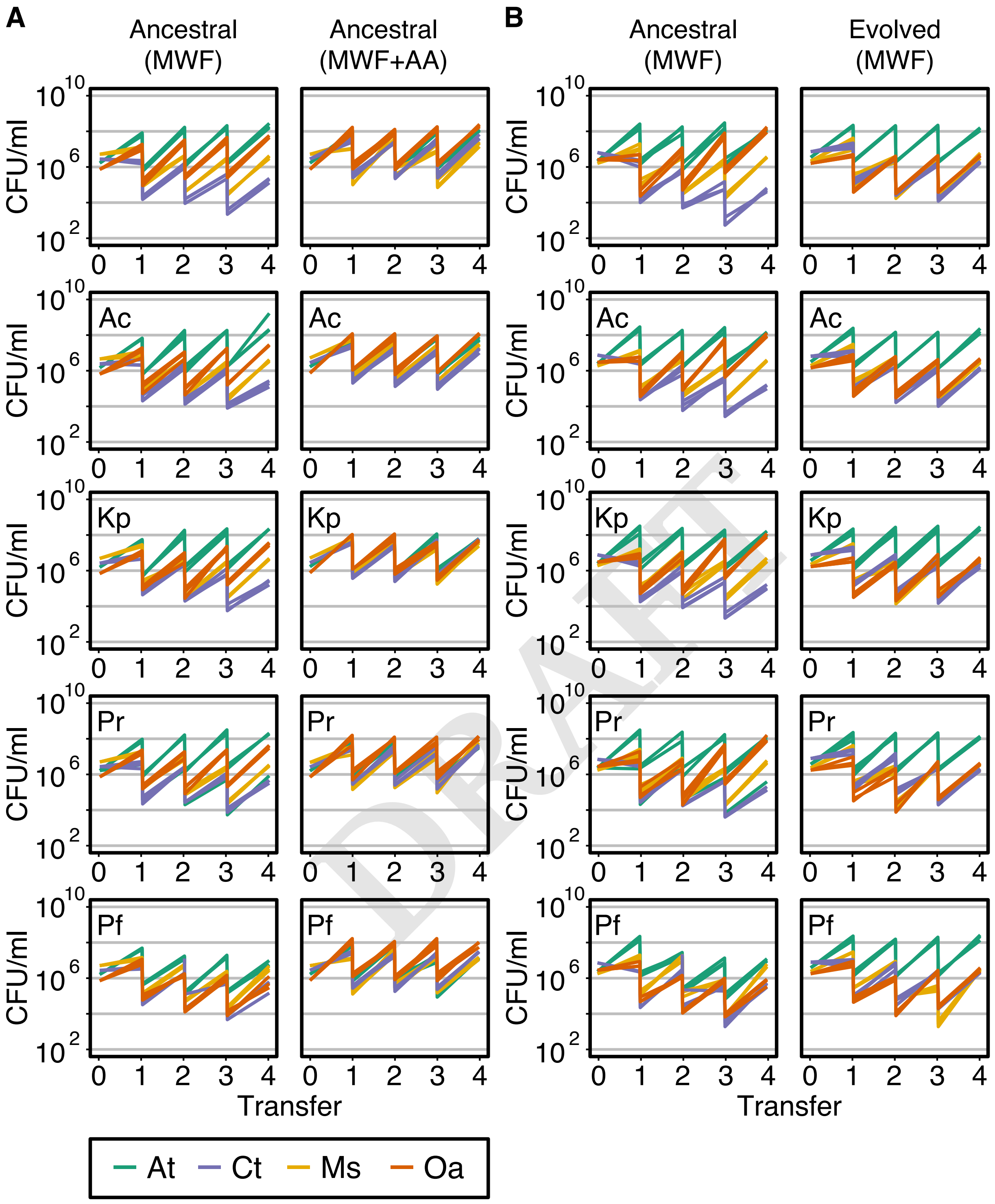


**Fig. S2.** Dynamics of the 4-species resident community. (A) Abundances of the ancestral resident species over the four transfers in MWF (left) or MWF+AA (right), quantified at first inoculation and before each transfer. Panel (B) shows ancestral species as a biological repeat (left) and co-evolved species (right). The four resident species were always grown together (resident community), with or without the inoculation of the invader. If present, the name of the invader is indicate in the top left of the plot. Invader dynamics are shown in the main text. The resident community was transferred every 7 days until transfer 4. At each transfer, the culture is diluted 100-fold. The experiments were done in parallel. Population sizes were quantified in Colony Forming Units (CFU) per milliliter at first inoculation and before each transfer. These data are summarized and compared in Fig. 3A, B and Fig. S7.

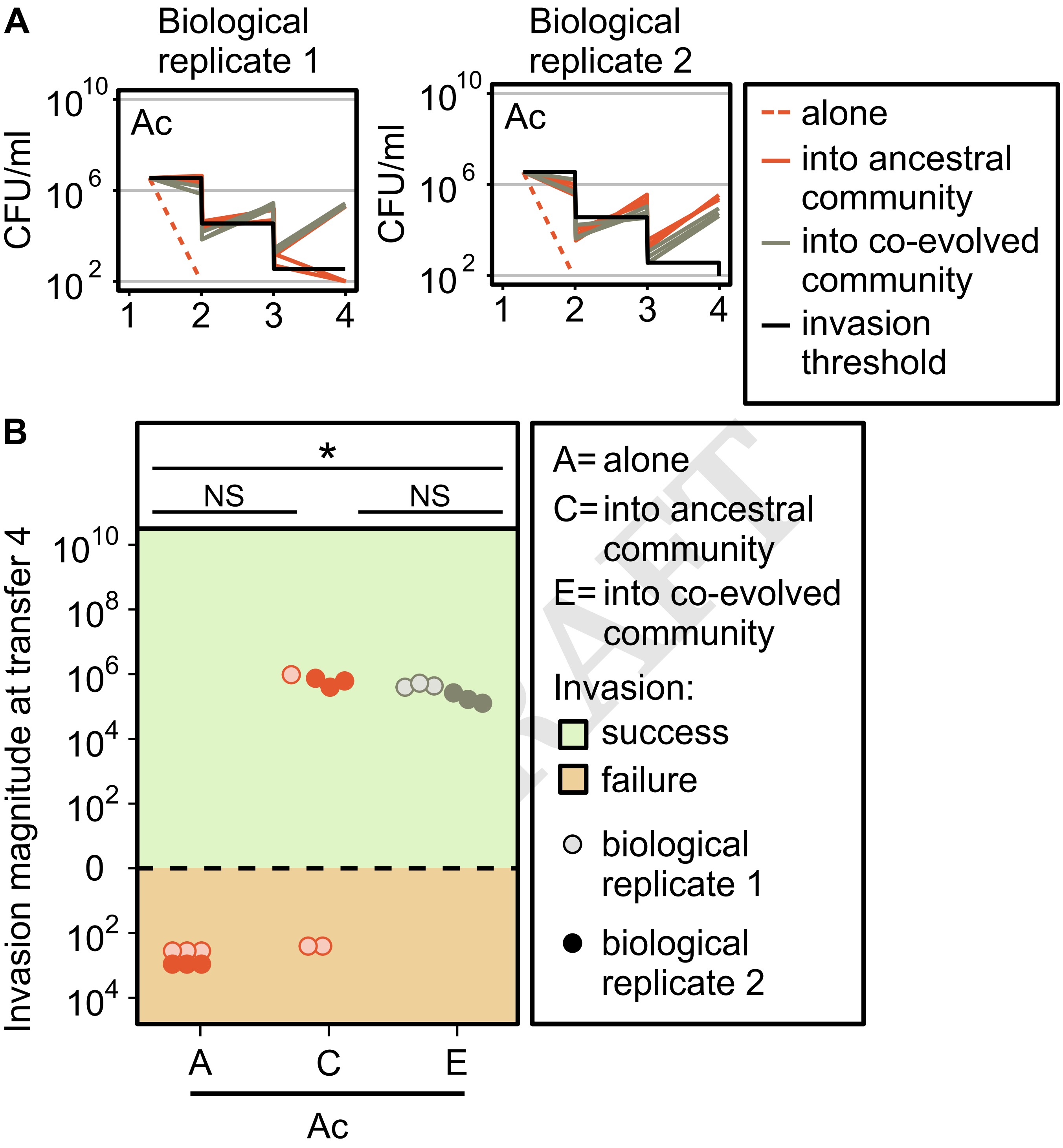


**Fig. S3.** Comparison of two biological replicates. (A) *A. caviae* (Ac) was grown alone, or inoculated into the ancestral or the co-evolved community in MWF. Populations were diluted 100-fold in fresh MWF every 7 days for a total of 4 transfers. Treatments in each biological replicate were all conducted in parallel. The black line represents the invasion threshold. (B) Invasion magnitude of two biological replicates of *A. caviae* (Ac) (population size at transfer 4 minus the invasion threshold). Positive or negative invasion magnitudes indicate successful (+) or failed (-) invasions, respectively. Population sizes were quantified in Colony Forming Units (CFU) per milliliter at first inoculation and before each transfer. Statistical significances are marked above the data points (Kruskal-Wallis, P-values: * $<$0.05, NS = not significant).

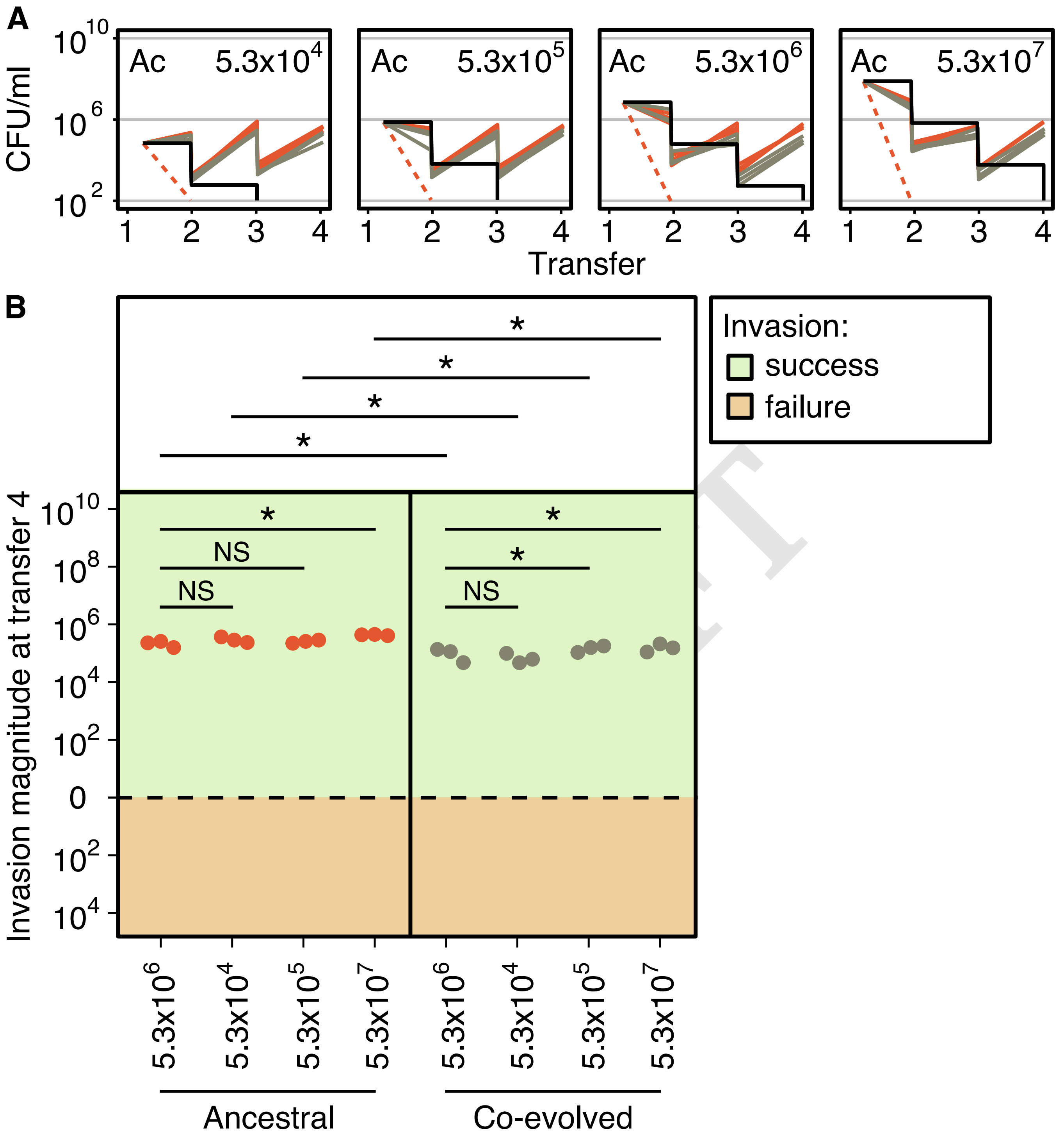


**Fig. S4.** Varying inoculum size of *A. caviae*. (A) *A. caviae* (Ac) was grown alone, or inoculated into the ancestral or the co-evolved community in MWF at four different population sizes. The inoculum size of *A. caviae* is indicated in the top right of the plot. Populations were diluted 100-fold in fresh MWF every 7 days for a total of 4 transfers. All conditions were conducted in parallel. The black line represents the invasion threshold. (B) Invasion magnitude of *A. caviae* (population size at transfer 4 minus the invasion threshold). Positive or negative invasion magnitudes indicate successful (+) or failed (-) invasions, respectively. Population sizes were quantified in Colony Forming Units (CFU) per milliliter at first inoculation and before each transfer. We compared the data points of the invader when inoculated at standard populations size (in general around 106, in this example is 5.3x106) with other treatments where the invader was diluted 100x, 10x or concentrated 10x before inoculation. Statistical significance are marked above the data points (Kruskal-Wallis, P-values: * $<$0.05, NS = not significant). Despite very different population sizes at invasion, all *A. caviae* populations end up at very similar abundances at transfer 4, which was consistently higher for the ancestral community compared to the evolved one.

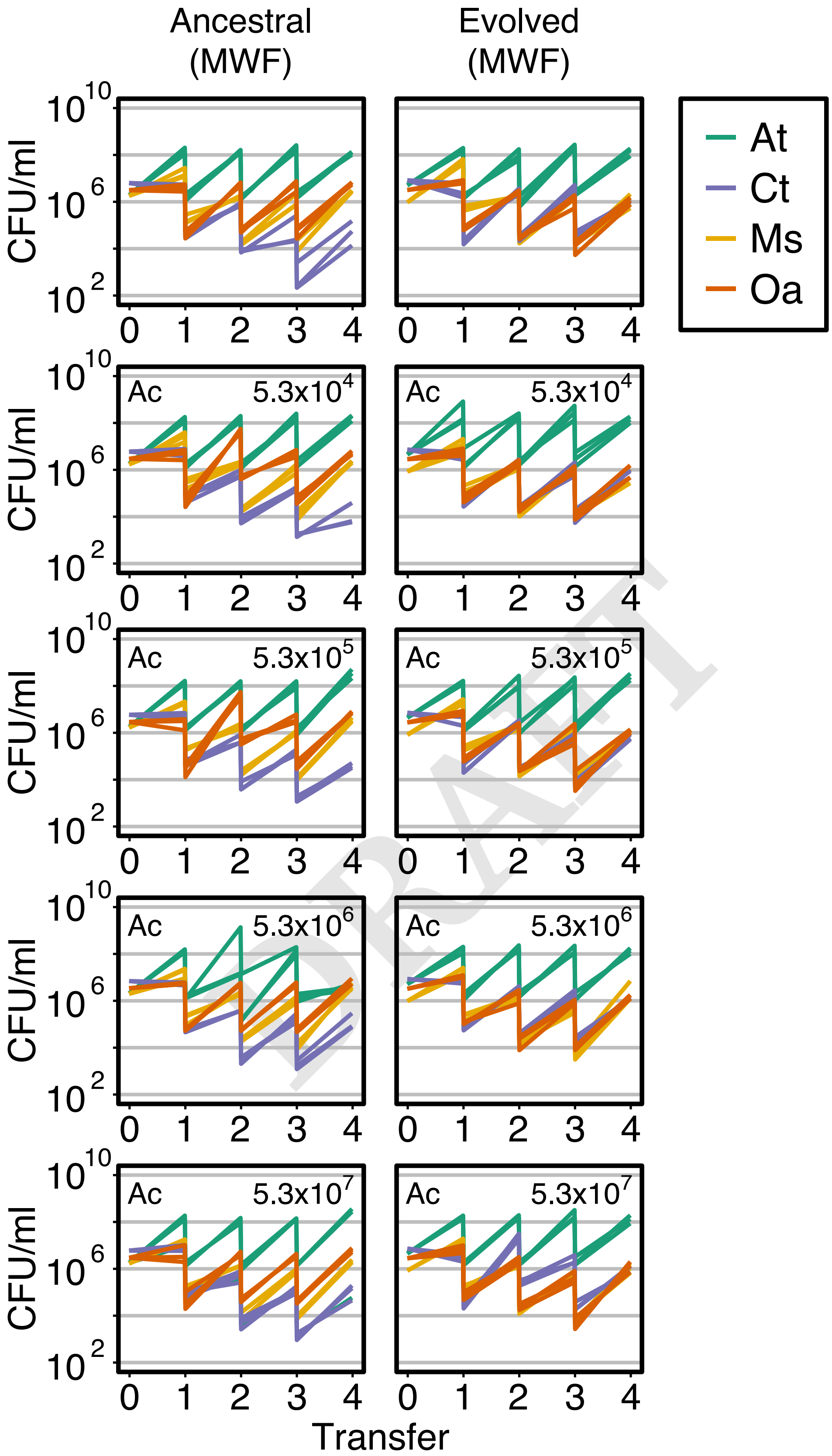


**Fig. S5.** Dynamics of the 4-species resident community by varying inoculum size of *A. caviae*. Abundances of the ancestral (left) or evolved (right) resident species over the four transfers in MWF, quantified at first inoculation and before each transfer. The four resident species were always grown together (resident community), with or without the inoculation of *A. caviae*. If present, the name of the invader is indicate in the top left of the plot. The inoculum size of *A. caviae* is indicated in the top right of the plot. The resident community was transferred every 7 days until transfer 4. At each transfer, the culture is diluted 100-fold. The experiments were done in parallel. Population sizes were quantified in Colony Forming Units (CFU) per milliliter at first inoculation and before each transfer.

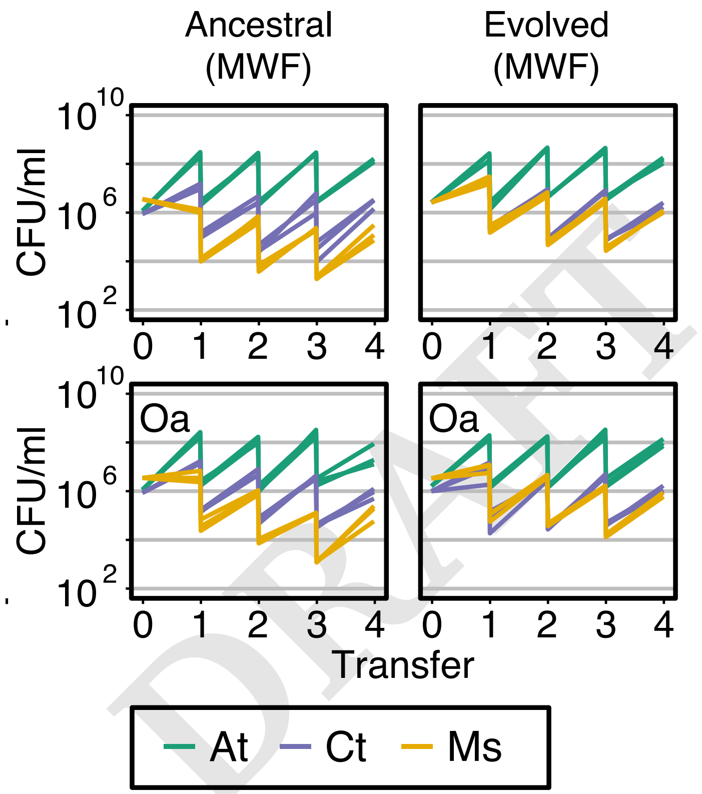


**Fig. S6.** Dynamics of the 3-species resident community. Abundances of the ancestral (left) or evolved (right) resident species over the four transfers in MWF, quantified at first inoculation and before each transfer. The three resident species were always grown together (resident community), with or without the inoculation of the invader (Oa). If present, the name of the invader is indicate in the top left of the plot. The resident community was transferred every 7 days until transfer 4. At each transfer, the culture is diluted 100-fold. The experiments were done in parallel. Population sizes were quantified in Colony Forming Units (CFU) per milliliter at first inoculation and before each transfer. These data are summarized and compared in Fig. 3C, D.

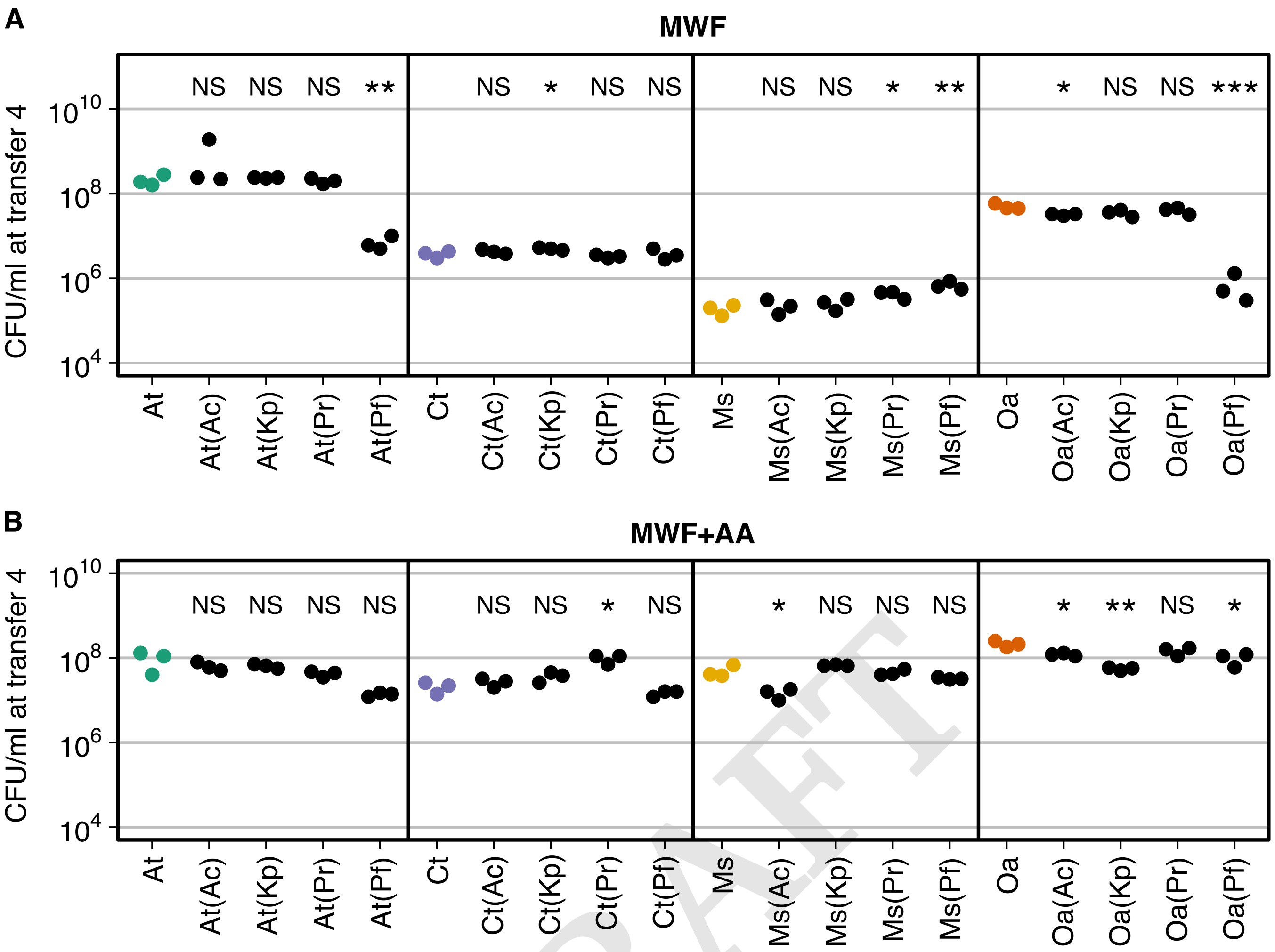


**Fig. S7. CFU/ml at transfer 4 of ancestral community members with or without invader in MWF or MWF+AA.** Each panel represents the total population size (CFU/mL) at transfer 4 of an ancestral resident species in MWF (A) or MWF+AA (B) when growing in the community of four with or without invaders. Species population sizes without any invader species are represented in colored dots and in black dots when an invader was added 48h after the first transfer (invader identity indicated in brackets). From left to right: *A. tumefaciens* (At), *C. testosteroni* (Ct), *M. saperdae* (Ms), *O. anthropi* (Oa), *A. caviae* (Ac), *K. pneumoniae* (Kp), *P. rettgeri* (Pr), *P. fulva* (Pf). We compared the data points of each species when invaded to the corresponding data points when co-cultured with other community members but without invasion. Statistical comparisons are marked above the co-culture data points (Kruskal-Wallis, P-values: * $<$0.05, **$<$0.01, NS = not significant). Panel (A) represents a biological replicate of Fig. 3A and shows similar results.

DRAFT

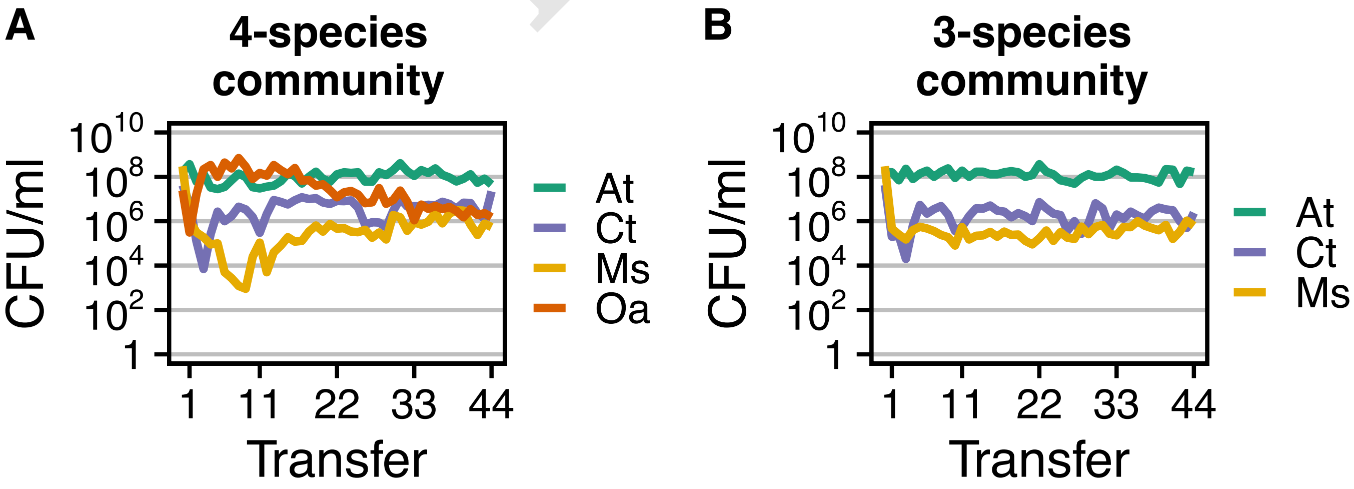


**Fig. S8.** Population sizes before each of 44 transfers. Experiments were started with each species assembled in co-cultures of four (*A. tumefaciens*, *C. testosteroni*, *M. saperdae* and *O. anthropi*) in panel (A) or three (*A. tumefaciens*, *C. testosteroni* and *M. saperdae*) in panel B. In each treatment, cells were inoculated into 5 microcosm replicates (culture tubes), of which we show one replicate here. To stimulate active growth, serial transfer was performed by diluting each culture 100-fold in fresh MWF once per week for 44 weeks. Before each transfer, cultures were quantified separately by selective plating.

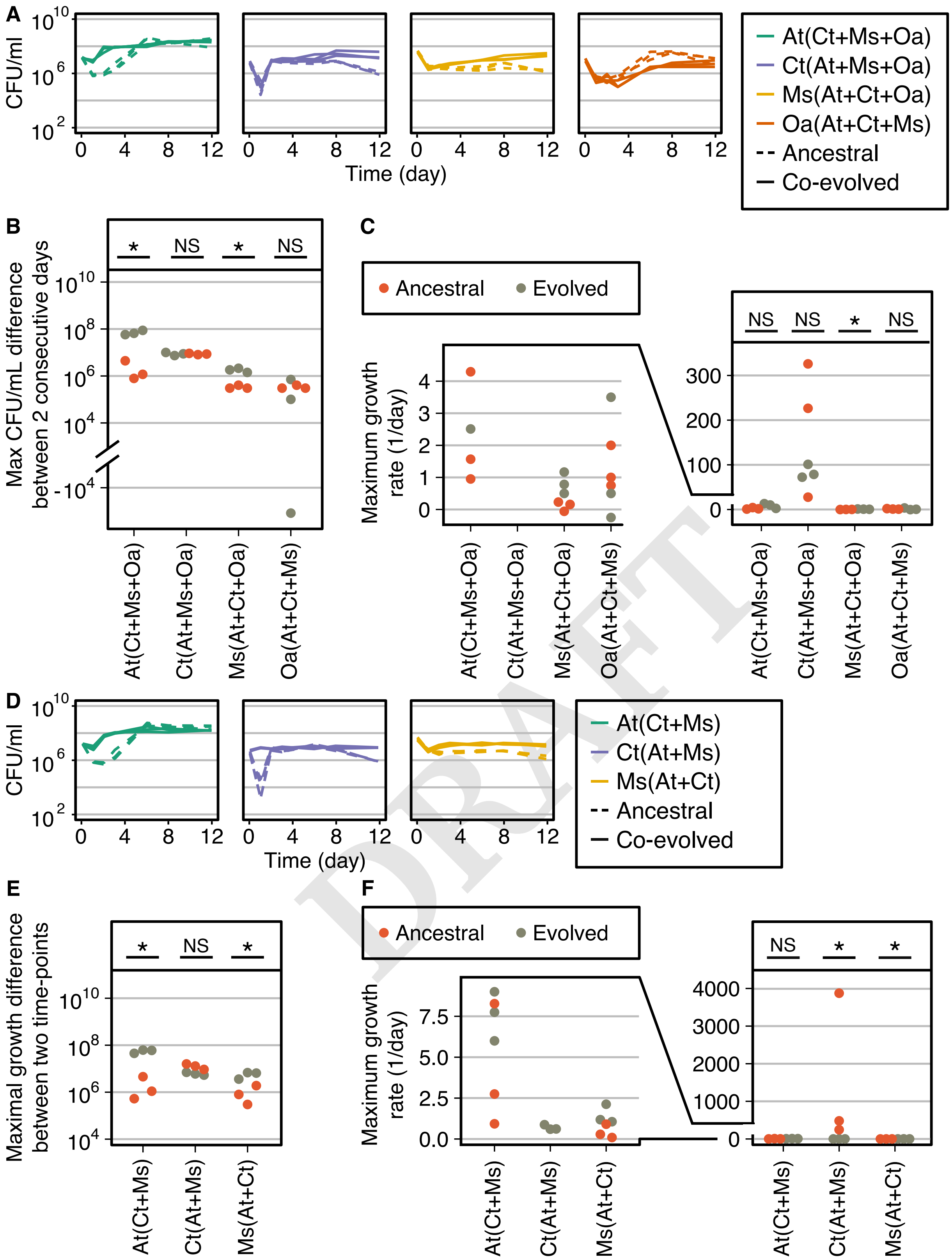

DRAFT

**Fig. S9. Growth of ancestral and co-evolved resident community members in MWF.** (A, D) Growth curves of the ancestral or co-evolved resident species, co-cultured with the other community members (co-culture partner indicated in brackets). (B, E) Each panel represents the maximal CFU/mL difference between two consecutive days of each species, both ancestral or co-evolved, co-cultured with the other community members (indicated in brackets). (C, F) Each panel represents the maximum growth rate (1/day) of each species, both ancestral or co-evolved, co-cultured with the other community members (indicated in brackets). The values are extrapolated from an additional experiment where each species, ancestral or evolved, was co-cultured with other resident members for 12 days. From left to right: *A. tumefaciens* (At), *C. testosteroni* (Ct), *M. saperdae* (Ms), *O. anthropi*. Statistical significance for comparisons of ancestral versus evolved is marked above the data points (Kruskal-Wallis, P-values: NS = not significant, * $<0.05$). Ancestral 3-species and 4-species community data points come from (47). Overall, the results indicate that co-evolved residents tend to grow to larger population sizes during the first days in the MWF, relative to their ancestors.

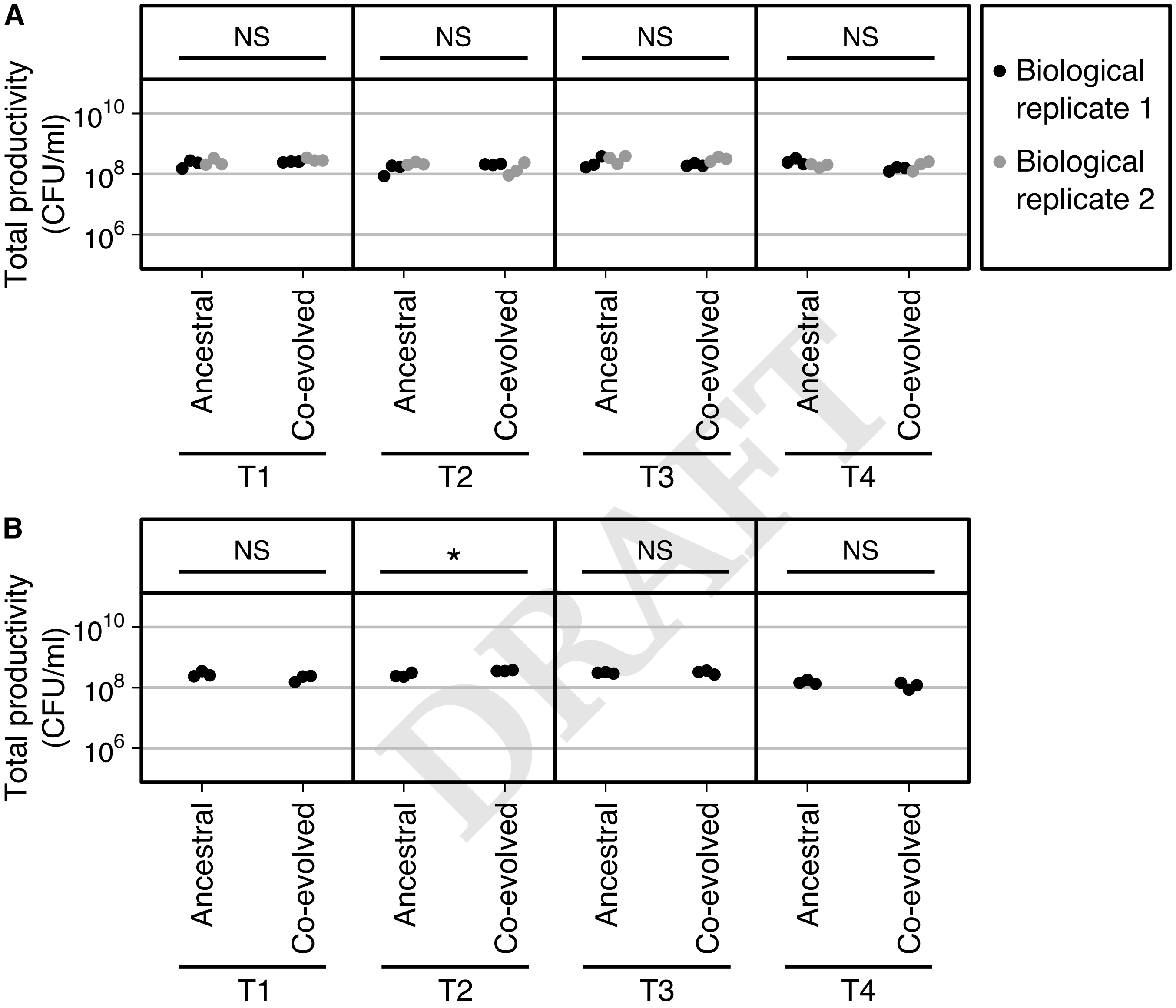


DRAFT

**Fig. S10.** Total community productivity. Comparison of the total CFU at the end of each transfer between ancestral and co-evolved communities, without invasion, in the co-evolved 4-species community (A), and 3-species community. Statistical significances are marked above the data points (Kruskal-Wallis, P-values: NS = not significant, * $<0.05$). Ancestral 3-species community data points come from (47).

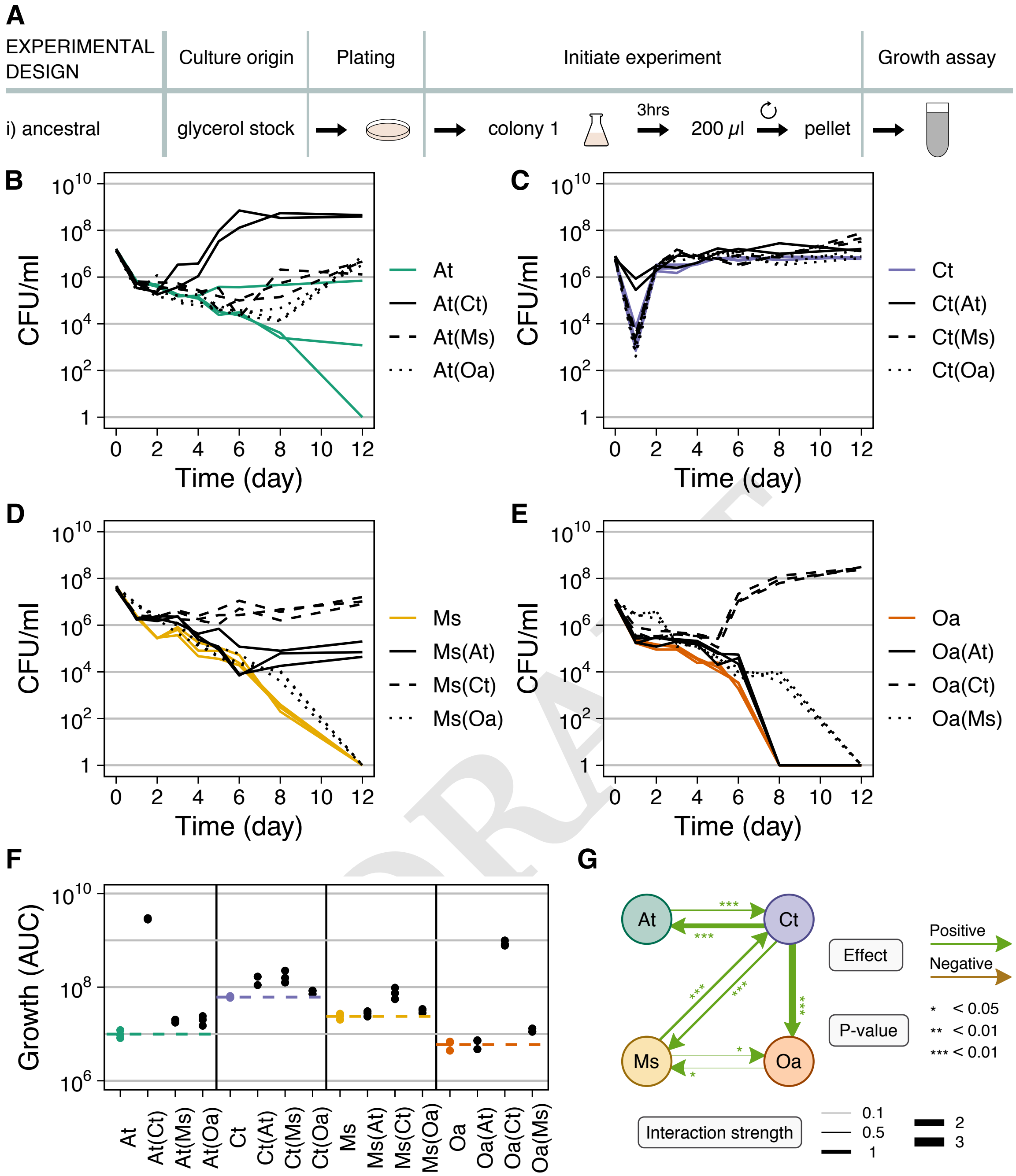


**Fig. S11.** Comparison of the ancestral bacterial community mono- and pairwise co-cultures, adapted from (47). (A) One colony of each ancestral species was randomly picked and grown alone for 3 hours to exponential phase, 200$\mu$L of each species mixed (if in a pair-wise co-culture, otherwise only 200$\mu$L were used), washed and resuspended in 30mL of MWF. (B-E) Population size quantified in Colony-Forming Units per milliliter over time for mono-cultures (in color) and pairwise co-cultures (in black; co-culture partner indicated in brackets). In the co-cultures, each species could be quantified separately by selective plating. Each panel shows the data for 1 species: (B) *A. tumefaciens* (At), (C) *C. testosteroni* (Ct), (D) *M. saperdae* (Ms) and (E) *O. anthropi* (Oa). (F) AUC calculated from data in B-E. Dashed lines indicate the mean of the mono-cultures, shown in color. (G) Pairwise interaction network. Positive/negative interactions indicate that the species at the end of an arrow grew significantly better/worse in the presence of the species at the beginning of the arrow. Arrow thickness represents interaction strength as the 10-fold change in the co-culture AUCs compared with mono-culture AUCs, i.e., by how many orders of magnitude a species changed the AUC of another. Dataset, statistical significance and interaction strengths data are provided in (47).

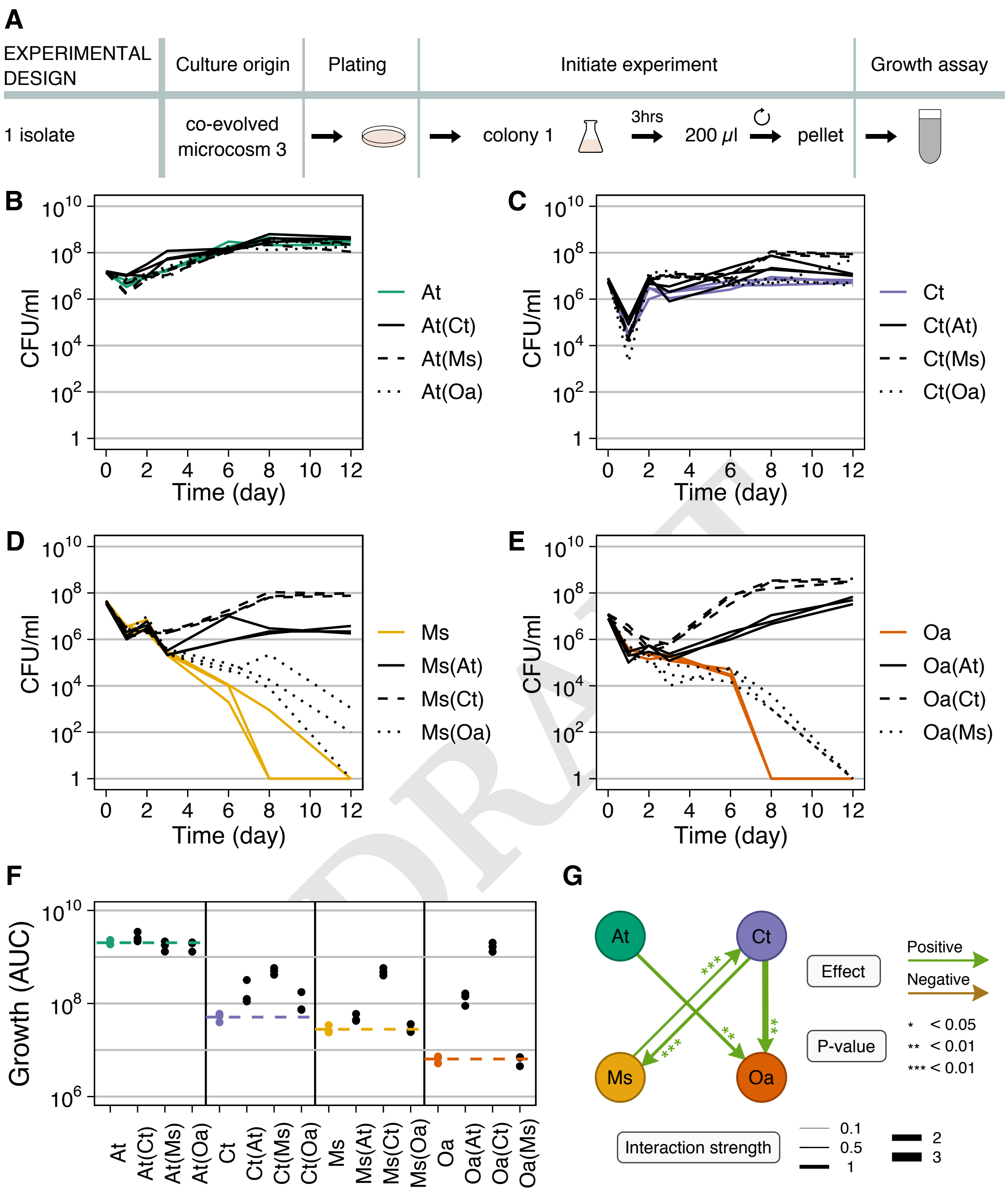


DRAFT

**Fig. S12.** Comparison of the 4-species co-evolved bacterial community mono- and pairwise co-cultures. (A) One colony of each species was randomly picked from the co-evolved community and grown alone for 3 hours to exponential phase, then washed, resuspended and mixed in equal proportions in MWF. (B-E) Population size quantified in colony-forming units per milliliter over time for mono-cultures (in color) and pairwise co-cultures (in black; co-culture partner indicated in brackets). In the co-cultures, each species could be quantified separately by selective plating. Each panel shows the data for 1 species: (B) *A. tumefaciens* (At), (C) *C. testosteroni* (Ct), (D) *M. saperdae* (Ms). (F) AUC calculated from data in B-D. Dashed lines indicate the mean of the mono-cultures, shown in color. (G) Pairwise interaction network. Positive/negative interactions indicate that the co-evolved species at the end of an arrow grew significantly better/worse in the presence of the co-evolved species at the beginning of the arrow. Arrow thickness represents interaction strength as the 10-fold change in the co-culture AUCs compared with mono-culture AUCs, i.e., by how many orders of magnitude a co-evolved species changed the AUC of another. Statistical significance and interaction strengths data are shown in Dataset S1.

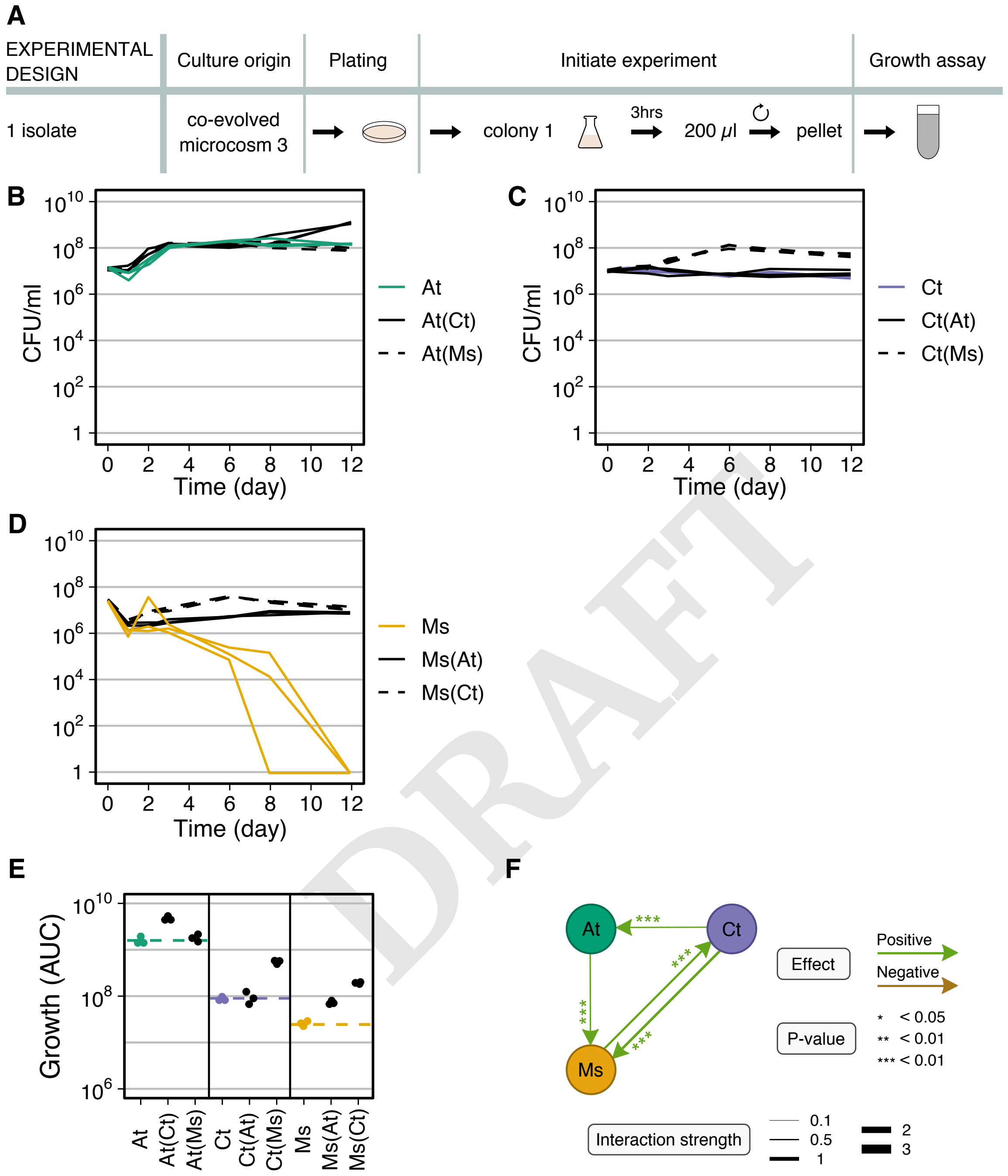


**Fig. S13.** Comparison of the 3-species co-evolved bacterial community mono- and pairwise co-cultures. (A) One co-evolved isolate of each species was randomly picked and grown alone 3 hours to exponential phase, then washed, resuspended and mixed in equal proportions in MWF. (B-D) Population size quantified in colony-forming units per milliliter over time for mono-cultures (in color) and pairwise co-cultures (in black; co-culture partner indicated in brackets). In the co-cultures, each species could be quantified separately by selective plating. Each panel shows the data for 1 species: (B) *A. tumefaciens* (At), (C) *C. testosteroni* (Ct), (D) *M. saperdae* (Ms). (E) AUC calculated using data in B-D. Dashed lines indicate the mean of the mono-cultures, shown in color. Dashed lines indicate the mean of the mono-cultures, shown in color. (F) Pairwise interaction network. Positive/negative interactions indicate that the co-evolved species at the end of an arrow grew significantly better/worse in the presence of the co-evolved species at the beginning of the arrow. Arrow thickness represents interaction strength as the 10-fold change in the co-culture AUCs compared with mono-culture AUCs, i.e., by how many orders of magnitude a co-evolved species changed the AUC of another. Statistical significance and interaction strengths data are shown in Dataset S1.

DRAFT
